# Experimental flowering order shapes priority effects in plant–pollinator interactions

**DOI:** 10.64898/2026.08.28.747010

**Authors:** Agostina Torres, Wei-Ling (Cherry) Chen, Janneke Hille Ris Lambers, Susan Waters

**Author notes:** Correspondence: Agostina Torres.

## Abstract

Climate change is disrupting life’s seasonal rhythms, altering the timing of key phenophases and reshaping how communities assemble. Beyond shifting flowering times, climate change can modify the extent of floral overlap and the sequence in which species bloom, generating novel assemblages with uncertain consequences for plant–pollinator interactions. Here, we ask whether flowering order generates priority effects in plant–pollinator communities, much like germination order does in plant communities. We tested how flowering order influences bee foraging behaviour and plant reproductive success in two co-flowering species, *Hypochaeris radicata* and *Campanula rotundifolia*, using a greenhouse experiment in which we manipulated the sequence of floral availability while allowing bees to forage repeatedly. We quantified changes in visit frequency, interspecific switches, handling time, and seed production. Our findings reveal priority effects in bee foraging that were strong enough to affect plant fitness: both species received more visits when flowering earlier than their co-occurring counterpart, and seed production declined when species flowered later. Overall, our results show that flowering order is an underappreciated driver of plant–pollinator interactions, suggesting that climate-driven phenological shifts could alter priority-effect dynamics with broader implications for community assembly. Key questions remain: How will climate-driven phenological shifts rearrange flowering sequences, and how will these priority effects emerge in more diverse communities in the wild? Our controlled experiment reveals strong flowering-order effects, underscoring the need to evaluate how widespread and impactful such dynamics are under accelerating climate change.

## Introduction

Climate change is disrupting life’s seasonal rhythms, altering the timing of key phenological events (Parmesan, 2007; Wolkovich et al., 2014). Among the most far-reaching consequences of these shifts are disruptions to plant–pollinator interactions, where altered flowering phenology can lead to mismatches that ripple through ecological networks, shaping the performance of co-occurring species (Kudo & Ida, 2013; Waters et al., 2020). Research has primarily focused on shifts in flowering timing, yet co-occurring species respond idiosyncratically to climatic cues, producing variation in the direction and magnitude of shifts (CaraDonna et al., 2014). As a result, climate change can alter not only when species flower, but also the extent of their overlap and the sequence in which they appear (e.g., Pareja-Bonilla et al., 2025), generating novel community assemblages with uncertain consequences for plant–pollinator interactions (Cleland & Wolkovich, 2024; Theobald et al., 2017).

Just as arrival order strongly influences plant interactions and community assembly (Fukami, 2015), flowering order may likewise structure plant–pollinator interactions through priority effects. Early-flowering species can gain a competitive advantage by attracting shared pollinators first, reducing visitation to later bloomers. Pollinator behaviour provides a mechanistic explanation: eusocial bees quickly learn flower-specific handling routines and, because switching between flower types is costly, they tend to persist on the same resource—a behaviour known as flower constancy (Chittka et al., 1999). On the bee side, this learning increases nectar and pollen intake efficiency (Waser, 1986), though benefits may diminish as resources become scarcer. On the plant side, such floral constancy can reduce interspecific pollen transfer and thus competition when flowering overlaps (Bolnick et al., 2003), though greater foraging efficiency may also shorten visits and lower per-visit pollen deposition (Mayberry et al., 2024). Early-flowering and floral constancy may interact to produce downstream effects by altering bee foraging patterns—changes that can ultimately affect plant reproductive success. Indeed, earlier-flowering plants can experience reduced competition for pollinators with co-flowering species (Forrest & Thomson, 2011; Kehrberger & Holzschuh, 2019), though they may also face shorter pollinator activity windows under cooler conditions or experience phenological mismatches (CaraDonna et al., 2014; Rafferty & Ives, 2011). In this context, priority effects arise because the order in which plant species begin flowering alters pollinator behavior, thereby influencing subsequent interactions and the reproductive success of later-flowering species. Regardless, whether shifts in flowering order indeed restructure pollinator interactions and ultimately alter plant fitness remains an open question.

Here, we assessed how flowering order influences bee foraging behavior and plant reproductive success for two co-flowering plant species that compete for pollinators (Waters et al., 2020). We asked whether pollinator visits and seed production increase when a species blooms before a competitor. Our goal was to compare pollinator visitation and seed production within each species across phenological treatments. Because both species are predominantly self-incompatible (Ortiz et al. 2006, Stevens et al. 2012, Waters et al. 2014), differences in seed production among treatments primarily reflect differences in pollination success. We conducted a greenhouse experiment with two prairie forbs, *Hypochaeris radicata* and *Campanula rotundifolia*, in which we manipulated the sequence of floral availability while allowing bees to forage repeatedly. We expected that flowering order influences: (1) pollinator visitation patterns, including visitation frequency, switches and handling time; and (2) plant reproductive success. Our study highlights how the temporal sequence of floral resource availability can structure bee foraging patterns and, in turn, plant fitness—offering insights into the temporal dimension of species interactions.

## Methods

We tested whether flowering order influenced bee foraging behavior and plant reproductive success using controlled floral arrays of two insect-pollinated prairie forbs, *Hypochaeris radicata* (introduced from Europe) and *Campanula rotundifolia* (native to North America). Both are perennial herbs that rely primarily on insect pollination for seed production and are predominantly self-incompatible (Ortiz et al. 2006, Stevens et al. 2012, Waters et al. 2014). *H. radicata* produces open, yellow composite flower heads that offer both nectar and pollen, whereas *C. rotundifolia* bears nodding, blue-violet bell-shaped flowers that primarily provide pollen. They co-occur in western Washington prairies, have overlapping summer flowering periods, and share a diverse assemblage of pollinators, including bumblebees (*Bombus* spp.), other Apidae, and Halictidae, creating opportunities for pollinator-mediated competition (Waters et al. 2020).

Western Washington prairies have declined to a small fraction of their historical extent due to land-use change, fire suppression, and nonnative species invasion, making them among the most endangered ecosystems in the Pacific Northwest (Noss et al., 1995). Ongoing climate change is expected to further affect these systems, not only through altered disturbance regimes and water availability (Bachelet et al., 2011), but also by shifting the phenology of native and nonnative plant species—potentially reshaping competitive and mutualistic interactions within these floral communities. Consistent with this expectation, experimental warming in Pacific Northwest prairies advanced first flowering by an average of 5.3 days per °C across eight focal species, although responses varied substantially among species (Reed et al. 2019). Likewise, *Campanula rotundifolia* flowered up to 20 days earlier than historical records in the northern Great Plains, illustrating the potential for substantial changes in flowering overlap over time (Dunnell and Travers 2011).

We manipulated the flowering sequence of *H. radicata* and *C. rotundifolia* across five consecutive days (Stages A-E) and offered bees different floral arrays in three treatments: *H. radicata* flowering before *C. rotundifolia*, *C. rotundifolia* flowering before *H. radicata*, and both species flowering simultaneously (Figure 1). The simultaneous treatment served as the baseline against which the effects of earlier or later flowering were evaluated. Each array consisted of 40 open flowers, with relative species abundances varying by treatment and day to simulate phenological scenarios (Figure 1). Experimental arrays were maintained at 40 open flowers by adding or removing entire pots and, when necessary, individual flowers from pots bearing multiple blooms. Although all flowers were at anthesis, time since anthesis was not standardized. Flowers were replaced daily to maintain standardized total and unpollinated floral displays throughout the experiment. This was not intended to mimic natural flower longevity, but rather to simulate shifts in flowering overlap resulting from changes in the timing of peak flowering between species.

**Figure 1.**
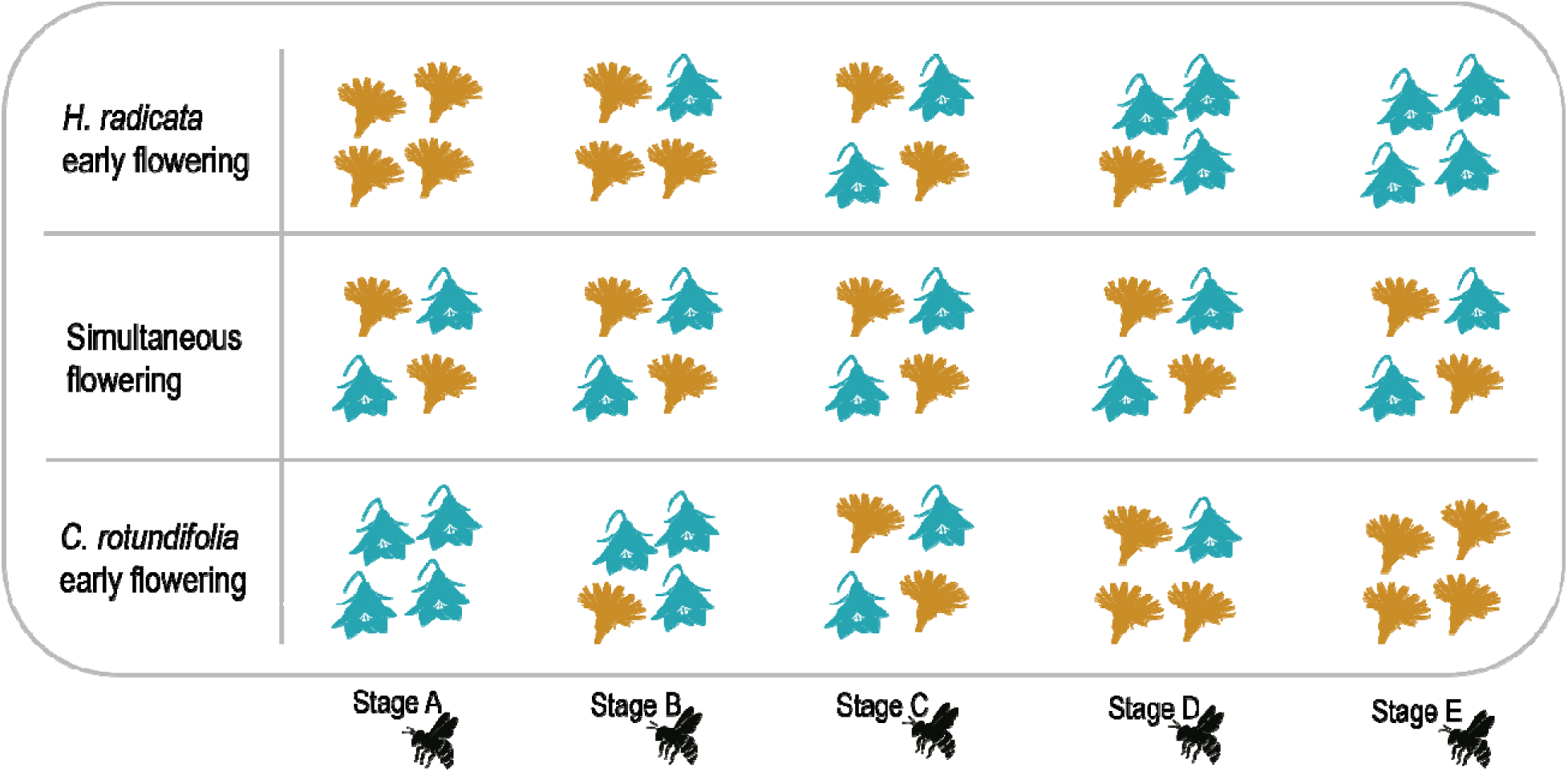
Experimental design. Each stage represents one of five consecutive days of the experiment (A to E), during which bumble bees foraged on floral arrays. Treatments simulated three flowering sequences using combinations of two plant species, *Hypochaeris radicata* (orange) and *Campanula rotundifolia* (blue), presented in 40-flower arrays. Here, each plant drawing represents 10 flowers. For early flowering, we started with all 40 flowers from the early-flowering species and, across five stages, progressively swapped them for the other species (40→30→20→10→0 flowers), mirroring the sequence when the other species arrived first. The simultaneous treatment stayed 50:50 (20 flowers of each) throughout all stages. A total of 30 unique *Bombus impatiens* individuals (10 from each of three colonies) were used across the entire experiment. Six unique bees per treatment were used in each experimental round in which that treatment was run (*Campanula early flowering* and *Hypochaeris early flowering* in both rounds; *Simultaneous flowering* in round 2 only). Each bee foraged once per day and was removed before the next bee was introduced. The foraging behavior of each bee was recorded during a complete foraging bout, defined as visiting at least 10% of the flowers with less than 30 seconds between visits. After foraging, visited flowers were bagged to measure seed production. For each treatment and species, seed number and weight were recorded.

For each scenario, we used six unique *Bombus impatiens* individuals from three colonies maintained under controlled conditions (24 °C daytime / 23.4 °C night, 12 h light:12 h dark; Table S1), which were sequentially released into a screened enclosure (0.65 m³). Bees were naïve at the start of each experimental scenario (Stage A), but the same individuals were subsequently reused on the following days, allowing them to retain any foraging experience acquired during the experiment (and thus, be affected by a behavioral priority effect if present). Each bee was allowed to forage once per day and removed before the next bee was introduced. After each daily foraging bout, marked bees were kept in separate containers in growth chambers between observation days. We recorded the number of visits to each species, the duration of foraging bouts, and the number of switches between species. A complete foraging bout was defined as bees visiting at least 10% of the flowers with fewer than 30 seconds between visits. After each bout, visited flowers were bagged to prevent further pollination and monitored for seed production (number and weight). Bagged flowers were replaced to restore each floral array to 40 open flowers before the next day’s observations.

### Data analyses

To evaluate bee foraging behaviour, we modelled visit frequency (visits flower^−1^ min^−1^), interspecific switches per bout, and handling time (bout duration/visit number) as functions of flowering order (early, simultaneous, late). We first analyzed visit frequency separately for each plant species using Tweedie GLMMs (log-link), which accommodate true zeros without pseudo-counts. We included flowering order treatment and experimental round as fixed effects, and bee ID (30 individuals) as a random intercept. Because visiting patterns can change across stages, we then fitted species-specific, stage-dependent GLMMs including stage, treatment, and experimental round as fixed effects, and bee ID as random effect. Stage-dependent models used Tweedie distribution (log-link) for visit frequency, a Gamma distribution for handling time and a negative binomial for switches.

To evaluate plant reproductive success, we modeled seed number (negative binomial) and weight (Tweedie) for each species separately using treatments and rounds as fixed effects. In addition, given that flowers were pooled from multiple individuals and their identity was not tracked, the hierarchical structure at the plant level could not be explicitly modelled. All models were implemented with *glmmTMB* (Brooks et al., 2017) in R 4.5.0 (R Core Team, 2025). Reported *p*-values for fixed effects are unadjusted Wald tests against the reference level. Estimated marginal means and additional pairwise contrasts were obtained with *emmeans* (Lenth et al., 2025), with the latter Tukey-adjusted for multiple comparisons. Model fit was assessed with DHARMa (Hartig, 2026).

## Results

Results show that bee behaviour is contingent on the flowering order. Both plant species had higher pollinator visits when flowering earlier than the co-occurring species (Figure 2A, C). In *Campanula rotundifolia*, visitation increased by 123% relative to simultaneous flowering, though this difference was not statistically robust, and by 184% relative to later flowering. Similarly, in *Hypochaeris radicata*, visitation was 141% higher compared to simultaneous flowering and 192% higher compared to later flowering (Figure 2A, C, Table S1). Visit frequency to early-flowering plants increased from Stage A to D despite declining relative abundance, whereas visit frequency under simultaneous flowering remained more stable over time (Figure 2B, D). This priority advantage was more pronounced and consistent across foraging stages for *Hypochaeris radicata*, where early-arriving plants received significantly higher visit frequencies from Stage B through D, while treatment differences across stages were comparatively modest for *Campanula rotundifolia,* with a visitation advantage emerging only by Stage D (Figure 2B, D, Table S2). Interspecific switches per bout did not differ among treatments or across stages (Figure S1A, Table S3). Estimated handling time (calculated as total bout duration divided by the number of visits to each species) was higher during the early–mid stages of the foraging sequence (Stages B–C) for both species when they flowered later (Figure S1B–C, Table S4), suggesting longer per-visit interaction times under delayed flowering.

**Figure 2.**
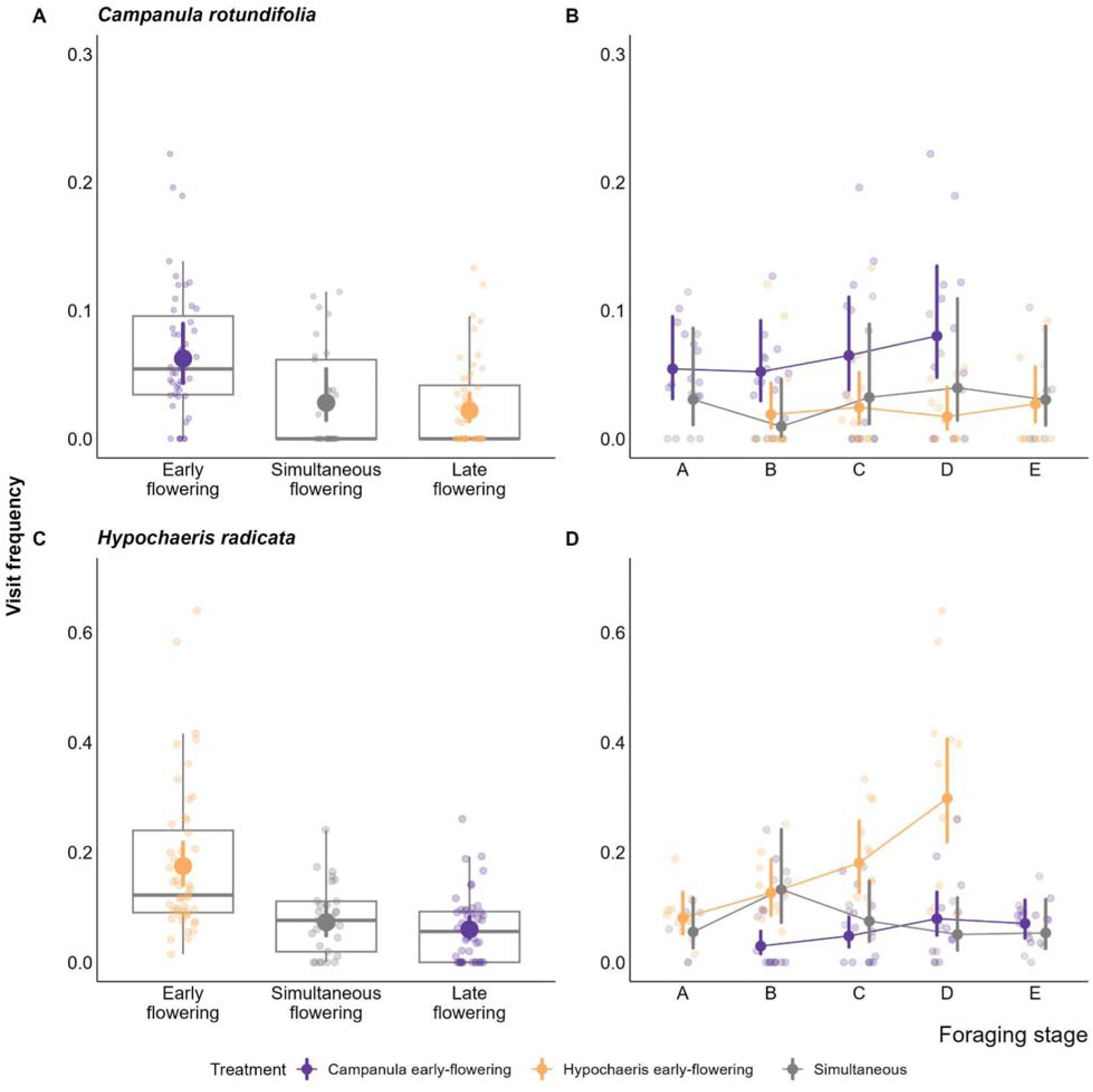
Effects of experimental flowering order on pollinator foraging behaviour. Visit frequency (visits flower ¹ min ¹) is shown for *Campanula rotundifolia* and *Hypochaeris radicata* across flowering-order treatments. Panels A and C show visit frequency across flowering-order treatments (early, simultaneous, late), averaged across foraging stages. Panels B and D illustrate the temporal progression of visit frequency across stages A–E within each treatment. Model predictions (coloured points with 95% CI) are shown together with raw data (boxplots in A and C; faint points in all panels).

Changes in bee foraging translated into plant reproductive output. Seed production was lower for both species when flowering late (Figure 3, Table S5 and S6). Both *Campanula* and *Hypochaeris* produced fewer seeds in the late-flowering treatment than in the early-flowering treatment (Figure 3A, B, Table S5). Our models estimate that *Campanula* produced 85% and *Hypochaeris* 73% fewer seeds when flowering late compared to flowering early. In addition, *Hypochaeris* seeds were ∼70% lighter when flowering late compared with early or simultaneous flowering (Figure 3D); whereas seed mass did not differ significantly among flowering-order treatments in *Campanula* (Figure 3C, Table S6).

**Figure 3.**
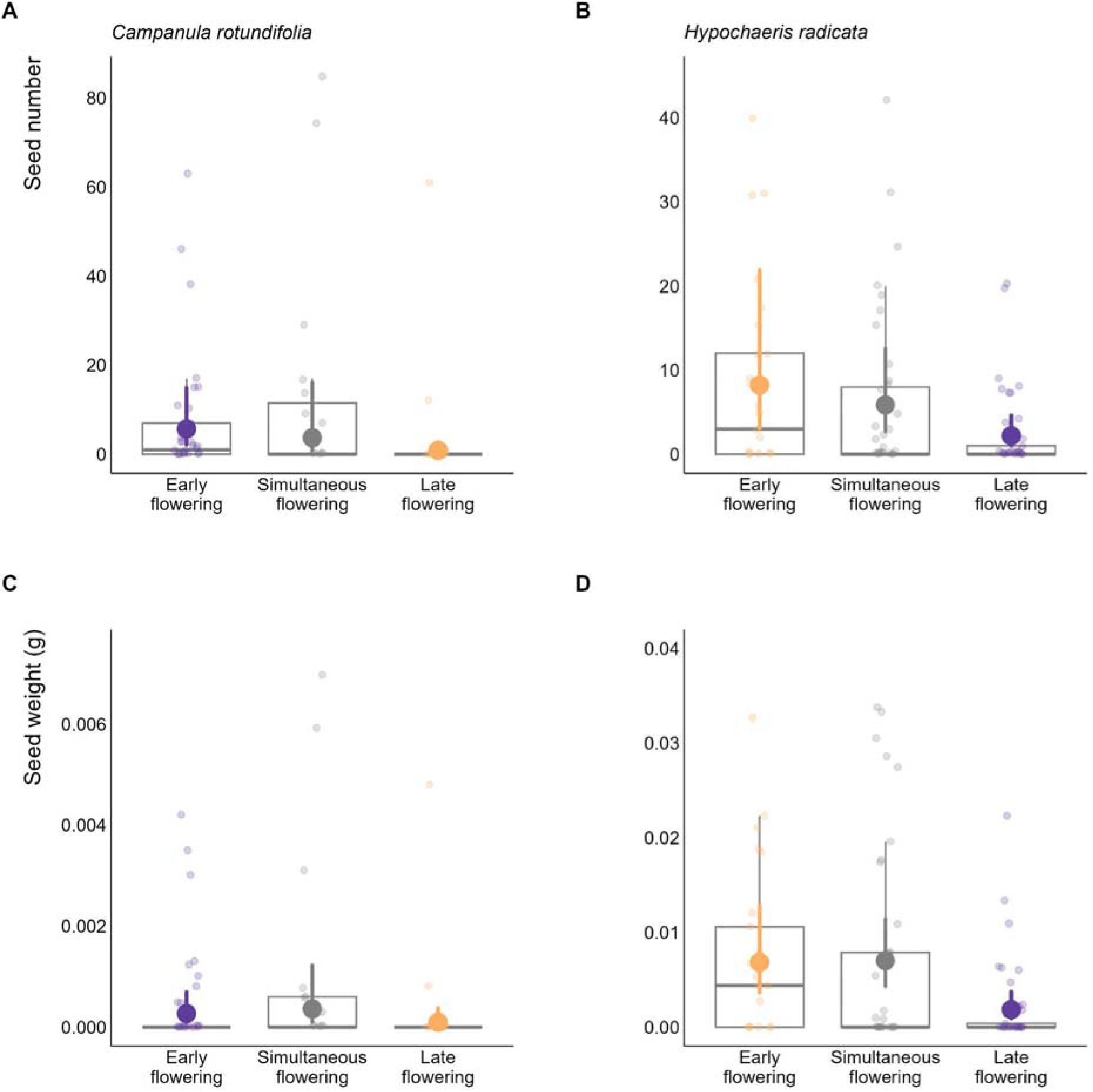
Effect of experimental flowering order on plant fitness. Seed number (A–B) and seed weight (g, C–D) produced by *Campanula rotundifolia* (purple) and *Hypochaeris radicata* (orange) across flowering order treatments (early, simultaneous, late). Colored dots with vertical bars (±95% CI) show model estimates; grey boxplots and faint points represent raw data.

## Discussion

Our findings reveal contingencies in bee foraging and plant fitness that depend on flowering order. Both focal species received more pollinator visits when flowering earlier than their co-occurring counterpart. Furthermore, changes in flowering order translated into plant fitness differences: seed production was lower when species flowered later, especially for *Hypochaeris*. Overall, our results reveal priority effects driven by shifts in flowering order, implying that climate-driven phenological changes in Washington prairies could alter priority effects dynamics, with potential consequences for community assembly.

Bee foraging behaviour varied with flowering order: visit frequency increased on the early-flowering species. In theory both early- and late-flowering species could benefit from reduced floral competition through temporal niche partitioning, but only early-flowering species showed increased visitation. One likely explanation is pollinator constancy—the tendency of bees to continue visiting the first species they learn to handle efficiently, reducing cognitive and energetic costs (Gegear & Laverty, 2005; Waser, 1986). Consistent with this, interspecific switching remained stable across treatments, even when the initially abundant species became rarer. While optimal foraging theory predicts that pollinators may adjust their foraging decisions as the relative abundance of floral resources changes (MacArthur and Pianka 1966, Pyke et al. 1977), we found little evidence for increased switching. This suggests that floral constancy or other behavioural constraints may have limited interspecific switching despite changes in floral abundance, at least in this system. Differences in floral morphology between *Campanula* and *Hypochaeris* may also have contributed to this pattern. The two species differ markedly in floral architecture and reward presentation which may increase the costs of switching between species. Such flower differences may reinforce floral constancy by favouring repeated visits to familiar flower types, thereby strengthening priority effects.

Handling-time patterns support the idea that pollinator constancy contributes to the higher visitation of early-flowering species: bees spent more time per visit on the later-flowering species during early–mid stages, suggesting higher handling costs when interacting with less familiar flowers (Laverty 1994; Grüter et al. 2011). Although nutritional needs can influence foraging decisions—bees typically balance pollen and nectar intake—this does not appear to override the priority effect in our system. Despite *Hypochaeris* providing both pollen and nectar, bees still concentrated their visitation on the species encountered first, indicating that priority effects can override immediate nutritional preferences. Together, these patterns suggest that the temporal order of floral availability can create historical contingencies in foraging trajectories that persist even when reward types are mixed.

Shifts in flowering order altered pollinator behavior in a way that translated into plant fitness. The advantage of flowering first—reflected in increased pollinator visitation and seed number for both species—indicates that even small changes in phenological sequence can modify, or even reverse, competitive hierarchies, underscoring the historical contingency of these interactions (Yin & Rudolf, 2024). Previous studies have similarly shown that shifts in flowering phenology can produce strong indirect effects on co-occurring species (Waters et al., 2020). In natural plant communities, species rarely flower in isolation and shifts in flowering order may therefore influence competitive and facilitative interactions among multiple co-flowering species. By altering which species are encountered first by pollinators, phenological changes could modify visitation patterns, reproductive success, and ultimately abundances, with consequences that extend beyond pairwise interactions. This may be particularly relevant in the context of biological invasions, given evidence that nonnative species tend to advance their flowering phenology more strongly than natives (Reeb et al., 2020; Wolkovich et al., 2013), potentially further increasing the advantage of those that flower earlier in the season. Our results suggest that, within the conditions of our experiment, flowering earlier benefited multiple components of reproductive output in the nonnative *Hypochaeris radicata*, affecting both seed number and seed weight. This species-specific response may, at least in part, reflect species-specific traits that modulate the strength of priority effects on fitness (van Steijn et al., 2025)— such as pollinator dependence or reproductive strategy. Pollinator-mediated interactions between plants can range from competitive to facilitative, implying that priority effects such as we found can have a wide range of effects.

Isolating ecological mechanisms often requires simplifying complexity—and our experimental study was no exception. A key strength of our approach was its simplicity and control. By using a robust design, we could isolate the effect of flowering order on bee foraging and plant fitness—something rarely possible in natural conditions. Holding floral abundance, species identity, pollinator availability, and environmental conditions constant, allowed us to isolate a mechanism often confounded in the field. Similar controlled approaches have been crucial for revealing priority effects in complex communities (e.g., Fukami et al., 2010; Vannette & Fukami, 2014). However, this level of control comes with limitations. Although our design allowed us to evaluate how flowering order influences pollinator visitation and seed production, it did not include monoculture controls, which would have provided a clearer baseline for interpreting how heterospecific interactions contribute to the observed effects of flowering order on visitation and reproductive output within each species. Our experiment simulated flowering order without accounting for the environmental cues that shape phenology in nature. In real communities, abiotic conditions, species interactions, and pollinator dynamics vary with flowering time. Likewise, keeping temperature constant meant we could not capture its potential effects on bee activity (CaraDonna et al., 2018) or on plant traits (de Manincor et al., 2023). While our design allowed us to test a specific mechanism under controlled circumstances, it cannot replicate the complexity of natural systems. Future studies should explore how priority effects in flowering interact with other ecological filters to shape community assembly.

Our findings highlight priority effects through flowering order as an underappreciated driver of plant–pollinator dynamics, echoing recent calls for broader recognition of priority effects across ecological contexts (Stroud et al., 2024). Several key questions remain open and particularly relevant to global change: How does climate-driven phenological change alter the flowering sequence among co-occurring species? How do changes to flowering order interact with flower abundance, overlap duration, and overlap timing to jointly shape priority effects? And finally, what are the broader consequences of altered priority effects at the community level? The patterns observed here point to potential long-term implications for species population dynamics, as well as direct and indirect effects that may cascade through interaction networks— including multilayer networks involving pollinators, seed dispersers, and seed predators. Temporal network approaches offer a promising way to quantify the downstream consequences of shifts in flowering order, helping reveal whether priority effects scale up in multispecies communities (Yin & Rudolf, 2024). Understanding how climate change will affect the prevalence and consequences of priority effects for community reassembly remains a key challenge.

## Acknowledgements

We thank Doug Ewing, UW Biology Greenhouse Manager, for invaluable assistance in designing and setting up the enclosures. This project was inspired by an earlier observational study conducted at Glacial Heritage Preserve. We also thank Marília Gaiarsa, Subject Editor at Oikos, and two anonymous reviewers for their constructive comments, which substantially improved the manuscript.

## Data availability statement

The data and code supporting the findings of this study are available in Dryad at DOI: 10.5061/dryad.31zcrjf2n.

---

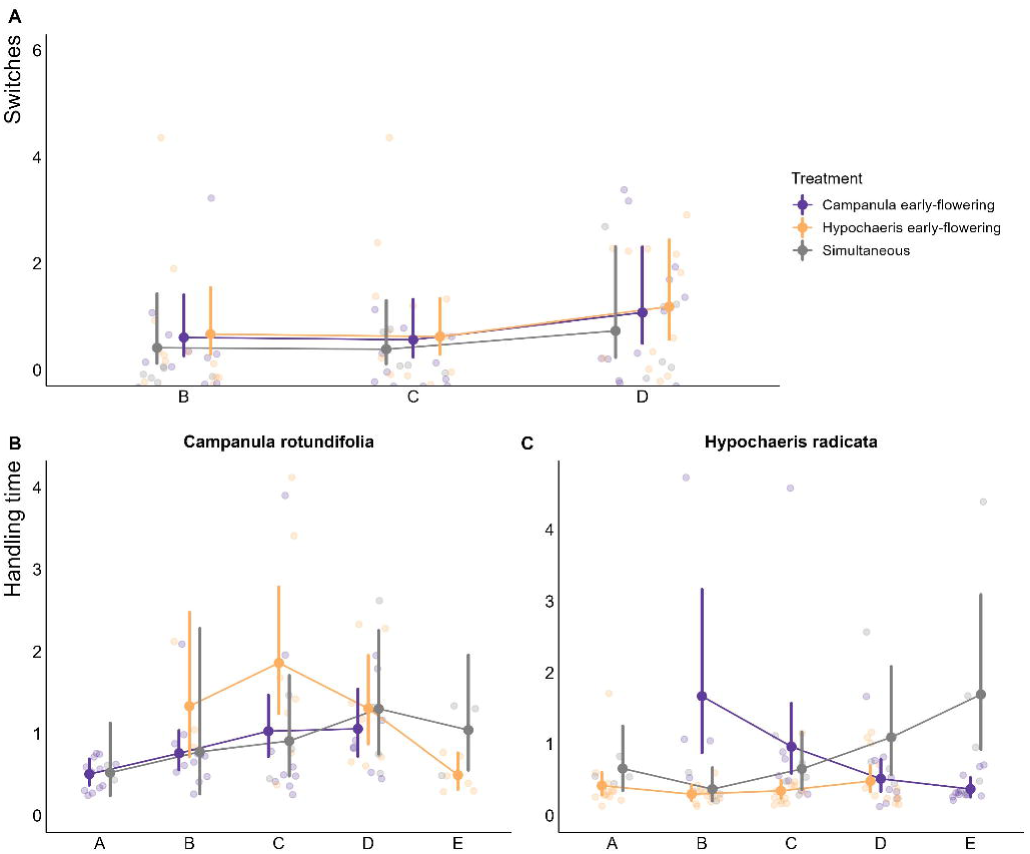

