## Supplementary material for "Experimental flowering order shapes priority effects in plant–pollinator interactions": S

**Figure S1**

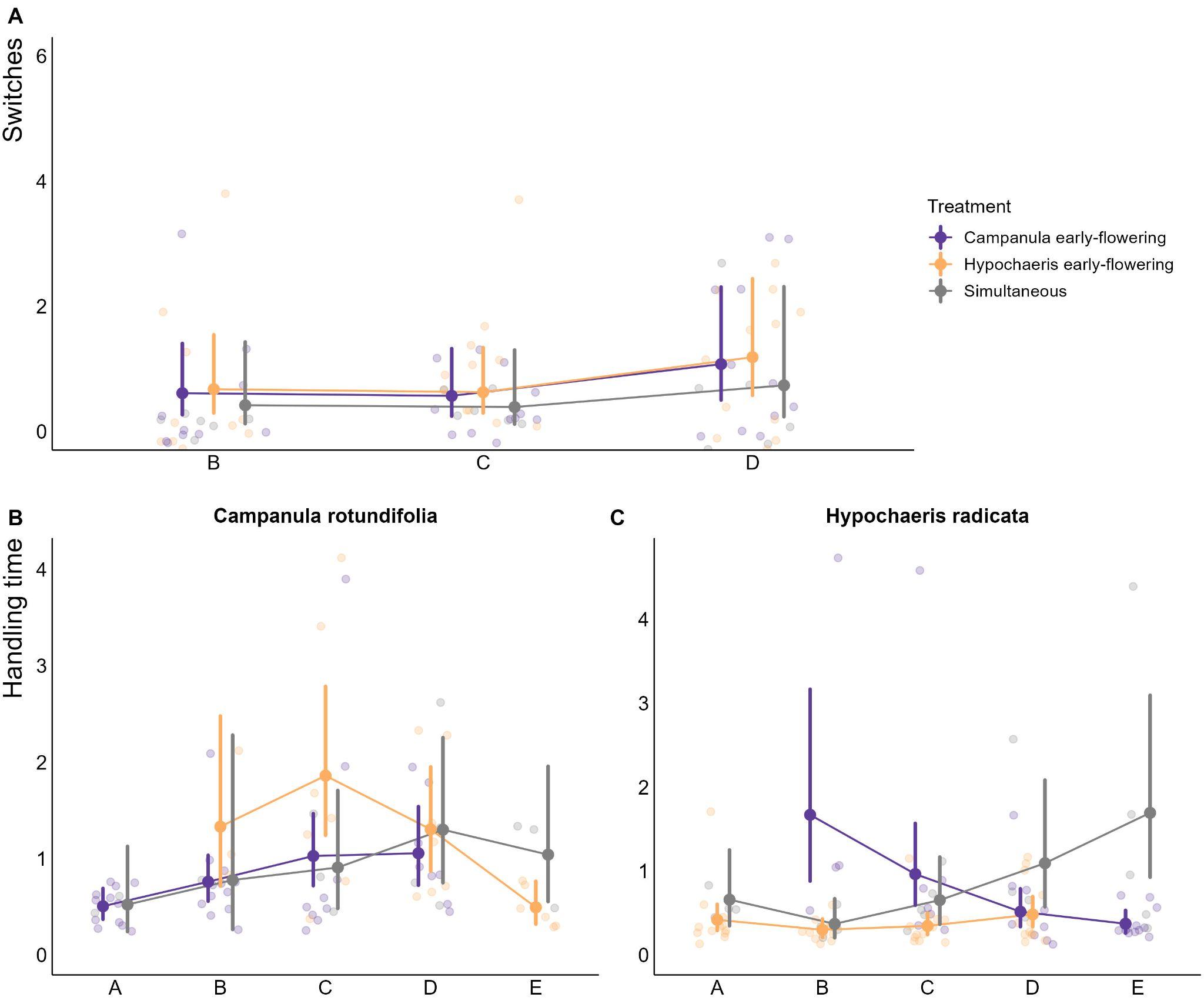

**Figure S1. Temporal changes in bee foraging behaviour across experiment stages and flowering-order treatments.** (A) Number of interspecific switches per foraging bout. (B–C) Handling time (bout duration / number of visits per bout) per flower for *Campanula rotundifolia* (left) and *Hypochaeris radicata* (right). Each panel shows model-estimated marginal means (±95% confidence intervals) for each treatment across stages, as obtained from the fitted GLMMs. Faded points represent individual observations. Colours indicate the different flowering-order treatments. One extreme value for the number of switches was excluded from the plot to improve visualization of the remaining data but was retained in all statistical analyses. For switches, we excluded stages A and E because switches cannot occur when only one species is flowering.

**Table S1. Visit frequency.** Results of two generalized linear mixed-effects models (GLMMs), one per plant species, testing the effect of flowering order and experimental round on bee visit frequency. Coefficients are presented on the log scale (Tweedie family, log link; n = 124 observations per species from 30 individual bees). Random intercept variance differed between plant species: for *Campanula*, the random effect of bee identity had a variance of 0.209 (SD = 0.458), whereas for *Hypochaeris* the variance was 0.034 (SD = 0.185).

|  | ***Campanula rotundifolia*** | | | | ***Hypochaeris radicata*** | | | |
| --- | --- | --- | --- | --- | --- | --- | --- | --- |
| **Predictive variable** | **Estimate** | **SE** | **z** | **p-value** | **Estimate** | **SE** | **z** | **p-value** |
| Intercept | -2.6830 | 0.233 | -11.52 | < 0.001 | -1.7880 | 0.147 | -12.16 | < 0.001 |
| Simultaneous flowering | -0.8010 | 0.378 | -2.12 | **0.034** | -0.8780 | 0.232 | -3.79 | **< 0.001** |
| Late flowering | -1.0430 | 0.298 | -3.51 | **< 0.001** | -1.0710 | 0.192 | -5.58 | **< 0.001** |
| Round | -0.1790 | 0.289 | -0.62 | 0.536 | 0.0940 | 0.189 | 0.50 | 0.618 |

**Table S2. Stage-level visit frequency.** Estimated coefficients (log scale) from GLMMs testing the effects of treatment, flowering stage, and their interaction on visit frequency to *Campanula rotundifolia* and *Hypochaeris radicata* separately. For *Campanula rotundifolia* (n = 200 observations from 30 bees) the random intercept for individual bee identity explained substantial variance (Var = 0.597, SD = 0.772). For *Hypochaeris radicata* (n = 201 observations from 30 bees) the random intercept for individual bee identity explained moderate variance (Var = 0.251, SD = 0.501). Some treatment × stage E interaction terms could not be estimated (NA), as flowering of the corresponding plant species had already ended by this stage, consistent with the experimental flowering-order design.

|  | ***Campanula rotundifolia*** | | | | ***Hypochaeris radicata*** | | | |
| --- | --- | --- | --- | --- | --- | --- | --- | --- |
| **Predictive variable** | **Estimate** | **SE** | **z** | **p-value** | **Estimate** | **SE** | **z** | **p-value** |
| Intercept | -2.81 | 0.32 | -8.88 | < 0.001 | -2.51 | 0.25 | -9.94 | < 0.001 |
| Late flowering | -0.70 | 0.82 | -0.85 | 0.394 | -0.10 | 0.58 | -0.17 | 0.866 |
| Simultaneous | -0.58 | 0.60 | -0.96 | 0.335 | -0.38 | 0.45 | -0.85 | 0.396 |
| Stage B | -0.04 | 0.36 | -0.12 | 0.907 | 0.45 | 0.27 | 1.68 | **0.093** |
| Stage C | 0.18 | 0.34 | 0.51 | 0.609 | 0.81 | 0.25 | 3.20 | **0.001** |
| Stage D | 0.39 | 0.34 | 1.15 | 0.251 | 1.31 | 0.24 | 5.50 | **< 0.001** |
| Stage E | 0.00 | 0.67 | 0.00 | 0.997 | -0.04 | 0.48 | -0.07 | 0.940 |
| Round | -0.20 | 0.29 | -0.69 | 0.491 | -0.01 | 0.20 | -0.06 | 0.950 |
| Late x B | -0.31 | 0.92 | -0.34 | 0.732 | -1.35 | 0.67 | -2.03 | **0.043** |
| Simultaneous x B | -1.11 | 0.96 | -1.16 | 0.245 | 0.42 | 0.50 | 0.85 | 0.395 |
| Late x C | -0.28 | 0.89 | -0.31 | 0.755 | -1.23 | 0.64 | -1.93 | **0.053** |
| Simultaneous x C | -0.12 | 0.73 | -0.16 | 0.871 | -0.50 | 0.52 | -0.98 | 0.329 |
| Late x D | -0.84 | 0.92 | -0.92 | 0.360 | -1.23 | 0.62 | -1.99 | **0.047** |
| Simultaneous x D | -0.13 | 0.72 | -0.17 | 0.863 | -1.39 | 0.57 | -2.46 | **0.014** |
| Late x E | NA | NA | NA | NA | NA | NA | NA | NA |
| Simultaneous x E | NA | NA | NA | NA | NA | NA | NA | NA |

**Table S3.** **Interspecific switches.** Estimated coefficients (log scale) from the negative binomial GLMM testing the effects of treatment, flowering stage, and round on the number of switches between plant species (n = 90 observations from 30 bees). The random intercept for individual bee identity was 0.1635 (SD = 0.4044).

| **Predictive variable** | **Estimate** | **SE** | **z** | **p-value** |
| --- | --- | --- | --- | --- |
| Intercept | -0.4680 | 0.708 | -0.66 | 0.508 |
| *Campanula* early-flowering | 0.3790 | 0.631 | 0.60 | 0.548 |
| *Hypochaeris* early-flowering | 0.4780 | 0.621 | 0.77 | 0.442 |
| Stage C | -0.0720 | 0.465 | -0.15 | 0.877 |
| Stage D | 0.5660 | 0.434 | 1.30 | 0.192 |
| Round | -0.8100 | 0.423 | -1.92 | **0.055** |

**Table S4.** **Stage-level handling time.** Estimated coefficients (log scale) from GLMMs testing the effects of treatment, flowering stage, and their interaction on handling time on *Campanula rotundifolia* and *Hypochaeris radicata* separately (Gamma family, log link). For *Campanula* (n = 77 observations from 26 bees), the random intercept for individual bee identity was estimated at the boundary of the parameter space (Var ≈ 0), indicating no detectable individual-level variability beyond what is explained by the fixed effects. For *Hypochaeris radicata* (n = 103 observations from 30 bees), the random intercept for individual bee identity explained modest variance (Var = 0.076, SD = 0.275). Some treatment × stage E interaction terms could not be estimated (NA), as flowering of the corresponding plant species had already ended by this stage, consistent with the experimental flowering-order design.

|  | ***Campanula rotundifolia*** | | | | ***Hypochaeris radicata*** | | | |
| --- | --- | --- | --- | --- | --- | --- | --- | --- |
| **Predictive variable** | **Estimate** | **SE** | **z** | **p-value** | **Estimate** | **SE** | **z** | **p-value** |
| Intercept | -0.681 | 0.181 | -3.76 | < 0.001 | -0.8100 | 0.214 | -3.79 | < 0.001 |
| Late flowering | -0.710 | 0.567 | -1.25 | 0.211 | -1.0600 | 0.482 | -2.20 | **0.028** |
| Simultaneous | 0.034 | 0.419 | 0.08 | 0.936 | 0.4530 | 0.373 | 1.21 | 0.225 |
| Stage B | 0.406 | 0.224 | 1.81 | **0.070** | -0.3330 | 0.236 | -1.41 | 0.158 |
| Stage C | 0.708 | 0.242 | 2.93 | **0.003** | -0.1890 | 0.238 | -0.79 | 0.427 |
| Stage D | 0.735 | 0.250 | 2.94 | **0.003** | 0.1430 | 0.242 | 0.59 | 0.555 |
| Stage E | 0.688 | 0.497 | 1.38 | 0.166 | 0.9430 | 0.405 | 2.32 | **0.020** |
| Round | 0.001 | 0.139 | 0.01 | 0.994 | -0.1180 | 0.179 | -0.66 | 0.509 |
| Late x B | 1.271 | 0.667 | 1.91 | **0.057** | 2.7750 | 0.590 | 4.70 | **< 0.001** |
| Simultaneous x B | -0.010 | 0.703 | -0.01 | 0.988 | -0.2440 | 0.462 | -0.53 | 0.598 |
| Late x C | 1.304 | 0.630 | 2.07 | **0.038** | 2.0820 | 0.550 | 3.79 | **< 0.001** |
| Simultaneous x C | -0.157 | 0.553 | -0.28 | 0.777 | 0.1800 | 0.457 | 0.39 | 0.693 |
| Late x D | 0.921 | 0.634 | 1.45 | 0.146 | 1.1250 | 0.539 | 2.09 | **0.037** |
| Simultaneous x D | 0.175 | 0.534 | 0.33 | 0.743 | 0.3630 | 0.481 | 0.75 | 0.451 |
| Late x E | NA | NA | NA | NA | NA | NA | NA | NA |
| Simultaneous x E | NA | NA | NA | NA | NA | NA | NA | NA |

**Table S5. Seed number models.** Estimated coefficients (log scale) from negative binomial GLMs testing the effects of flowering order and round on seed number for *Campanula rotundifolia* and *Hypochaeris radicata* separately. For *Campanula rotundifolia* (n = 86 observations), seed number was significantly lower under late flowering compared to early flowering, and in round 2 compared to round 1. For *Hypochaeris radicata* (n = 100 observations), seed number was also significantly lower under late flowering compared to early flowering, but did not differ significantly between rounds.

|  | ***Campanula rotundifolia*** | | | | ***Hypochaeris radicata*** | | | |
| --- | --- | --- | --- | --- | --- | --- | --- | --- |
| **Predictive variable** | **Estimate** | **SE** | **z** | **p-value** | **Estimate** | **SE** | **z** | **p-value** |
| Intercept | 2.95 | 0.72 | 4.07 | < 0.001 | 2.15 | 0.48 | 4.51 | < 0.001 |
| Simultaneous flowering | -0.43 | 0.96 | -0.44 | 0.657 | -0.34 | 0.62 | -0.55 | 0.585 |
| Late flowering | -1.88 | 0.8 | -2.36 | **0.018** | -1.32 | 0.64 | -2.07 | **0.038** |
| Round | -2.42 | 0.8 | -3.04 | **0.002** | -0.09 | 0.5 | -0.18 | 0.857 |

**Table S6. Seed weight models.** Estimated coefficients (log scale) from Tweedie GLMs testing the effects of flowering order and round on seed weight for *Campanula rotundifolia* and *Hypochaeris radicata* separately. For *Campanula rotundifolia* (n = 85 observations), seed weight did not differ significantly by flowering order but was significantly lower in round 2 compared to round 1. For *Hypochaeris radicata* (n = 99 observations), seed weight was significantly lower under late flowering compared to early flowering but did not differ significantly between rounds.

|  | ***Campanula rotundifolia*** | | | | ***Hypochaeris radicata*** | | | |
| --- | --- | --- | --- | --- | --- | --- | --- | --- |
| **Predictive variable** | **Estimate** | **SE** | **z** | **p-value** | **Estimate** | **SE** | **z** | **p-value** |
| Intercept | -7.2 | 0.47 | -15.42 | < 0.001 | -4.94 | 0.32 | -15.54 | < 0.001 |
| Simultaneous flowering | 0.29 | 0.65 | 0.45 | 0.656 | 0.03 | 0.41 | 0.06 | 0.950 |
| Late flowering | -1.04 | 0.77 | -1.35 | 0.177 | -1.3 | 0.47 | -2.76 | **0.006** |
| Round | -2.04 | 0.86 | -2.36 | **0.018** | -0.08 | 0.36 | -0.24 | 0.813 |
